# Comprehensive Analysis of ^177^Lu-lilotomab Satetraxetan in Lymphoma Cell Lines: Implications for Precision Radioimmunotherapy and Combination Schemes

**DOI:** 10.1101/2024.05.30.596390

**Authors:** Sebastian Patzke, Luciano Cascione, Katrine B Melhus, Nicolas Munz, Alberto J. Arribas, Eugenio Gaudio, Roman Generalov, Ada HV Repetto-Llamazares, Jostein Dahle, Francesco Bertoni

## Abstract

^177^Lu-lilotomab satetraxetan (Betalutin) is an anti-CD37 radioimmunoconjugate evaluated as single administration therapy for the treatment of patients with relapsed/refractory follicular lymphoma or diffuse large B-cell lymphoma (DLBCL). ^177^Lu-lilotomab satetraxetan treatment is well-tolerated and shows consistent activity in most of the patients evaluated so far. Herein, we investigated the activity of ^177^Lu-lilotomab satetraxetan in a panel of 55 lymphoma cell lines of B and T cell origin. CD37-targeted radioimmunotherapy was more effective in CD37-positive B-cell lymphomas (n=46) than negative CD37 negative T-cell lymphomas (n=9). Focusing on DLBCL cell lines, mutations such as *BCL2* or *MYC* translocations were not correlated to sensitivity. However, *BCL2* expression was higher in resistant than sensitive GCB-DLBCL cell lines, and the addition of the BCL2 inhibitor venetoclax showed synergism when added to the radioimmunoconjugate. Finally, the pattern of activity of ^177^Lu-lilotomab satetraxetan differed from what was achieved with a CD37-targeting antibody-drug conjugate or with R-CHOP, indicating the potential benefit of the beta-emitter payload. In conclusion, this systematic analysis of the responsiveness of lymphoma cell lines to CD37-targeting radioimmunotherapy consolidated ^177^Lu-lilotomab satetraxetan as a promising compound for the treatment of CD37 positive malignancies and identified candidate biomarkers and co-targets to detect and overcome cancer cell-intrinsic resistance mechanisms.

## Introduction

Lymphomas are among the most radiosensitive tumors ^1^. Labeling monoclonal antibodies targeting molecules on the surface of the lymphoma cells with radionuclides allows selective delivery of ionizing radiations to tumor cells ^12,3^. Based on the results of phase III randomized trials, two first-generation radiolabeled murine anti-CD20 monoclonal antibodies ^90^Y ibritumomab tiuxetan (Zevalin) and ^131^I tositumomab (Bexxar) were approved by the Food and Drug Administration (FDA) for relapsed or refractory low grade follicular or transformed B-cell lymphoma and relapsed or refractory CD20 positive follicular lymphoma, with and without transformation, respectively ^1,4,5^. Despite their strong anti-tumor activity, these radioimmunoconjugates have not been implemented in clinical practice, as testified by the withdrawal of ^131^I tositumomab in 2014 from the market for business reasons ^1^.

CD37 is widely expressed on mature normal and neoplastic B-cells ^6-10^, making it a potential therapeutic target. Indeed, one of the first demonstrations of the anti-tumor activity of radioimmunoconjugates in the clinical setting was with the anti-CD37 ^131^I MB-1 antibody given at high doses with autologous bone marrow support ^11^. CD37 has also been used as the target of antibody-drug conjugates (ADCs) ^6,7,12-14^. Lutetium-177 (^177^Lu) lilotomab satetraxetan (Betalutin) is a CD37-targeting radioimmunoconjugate obtained by conjugating the murine anti-CD37 IgG1 lilotomab with the bi-functional chelator p-SCN-benzyl-DOTA (satetraxetan) that chelates ^177^Lu^3+ 15^. Lutetium-177 is a β-emitting nuclide, a class of radionuclides suitable to treat lymphomas, with superior properties in availability, radiochemistry, and half-life ^3,15. 177^Lu-lilotomab satetraxetan has shown preclinical activity against small panels of lymphoma cell lines, including Burkitt lymphoma cell lines and activated B-cell-like (ABC) diffuse large B-cell lymphomas (DLBCL) ^15-19^. Clinical activity as a single agent has also been observed in indolent and aggressive lymphomas ^20,21^. An overall response rate of 70%, with 32% of complete responses, has been reported in 37 patients with second-line relapsed or refractory follicular lymphoma ^20^. The most frequent adverse events were reversible grade 3/4 neutropenia (32%), thrombocytopenia (26%), and grade 1/2 nausea (16%) ^20^. However, when ^177^Lu-lilotomab satetraxetan was tested in 3^rd^ line relapsed and CD20 refractory follicular lymphoma patients, the overall response rate dropped to 39 %, which was not considered competitive enough, and further development for ^177^Lu-lilotomab satetraxetan was discontinued (https://mb.cision.com/Main/9819/3752237/1990754.pdf).

Here, we present extensive data exploring the anti-tumor activity of ^177^Lu-lilotomab satetraxetan in 55 well-characterized lymphoma cell lines, focusing on the expression of its target and the identification of possible biomarkers and active combination partners.

## Materials and methods

H2>Cell lines

Cell lines were grown in the recommended medium supplemented with Glutamax (Gibco, Paisley, UK), 15% heat-inactivated fetal calf serum (Gibco), and 1% penicillin/streptomycin (Gibco) in a humid atmosphere with 95% air/5% CO_2_ and maintained in exponential growth phase through sub-culturing every 2-4 days.

### In vitro screen of ^177^Lu-lilotomab satetraxetan activity against lymphoma cell lines

Two replicates of each cell line were incubated in 96-deep-well plates with 12 different concentrations of ^177^Lu-lilotomab satetraxetan (obtained radiolabeling lilotomab satetraxetan with ^177^Lu), ranging from 0 to 20 μg/mL for 144 hours, as previously described ^19^. Data were analyzed by non-linear regression using GraphPad Prism version 7 to determine IC50 values (concentration at which the number of cells is half the number of control cells) and dose-response data compared across cell lines by clustering analysis ^19^.

### Data mining

TP53, BCL2, and MYC status had been previously determined in ^22^. CD37 RNA expression values were extracted from the datasets GSE94669 ^23^, previously obtained using a microarray-based technology (Illumina HT-12 arrays), and GSE221770 ^24^, previously produced via total-RNA-Seq. Differences in IC50 values among lymphoma subtypes were calculated using the Mann-Whitney test. P values of 0.05 or less defined statistical significance.

Differential gene expression analysis between groups of cell lines was performed using the voom/limma ^25^ R package on targeted RNA-seq data obtained with the HTG EdgeSeq Oncology Biomarker panel and included in dataset GSE94669 ^23^.

### Combinatorial drug treatment

Cells were seeded into deep-well 96 well plates at optimal cell concentrations. The small molecule was dispensed into the wells using a Tecan D300e digital drug dispenser at concentrations corresponding to IC25, IC50, and IC75 in quadruplets and three different sectors of the plate (that later would correspond to three different doses of ^177^Lu-lilotomab satetraxetan), gently mixed on a plate mixer and incubated for 1 hour in a humid atmosphere with 95% air/5% CO_2_. ^177^Lu-lilotomab satetraxetan was then co-dispensed to two plate sectors at 0.5 μg/mL and 1.0 μg/mL (final concentration), respectively. The third sector of the plate was left for control (0.0 μg/mL). A series of preliminary tests confirmed that the γ activity (low energy, co-originating from ^177^Lu decay) adjacent wells with the above-stated doses did not induce any noticeable cell toxicity within the time course of the experiment (data not shown). Plates were incubated for 18 hours, and then cells were washed three times with 0,9 mL of pre-warmed PBS. After washing, the cell pellets were reconstituted in the original volume of the pre-warmed cell growth medium. Cells were transferred to a new white flat bottom plate (75 μL/well), and a necessary amount of inhibitor was dispensed to restore the inhibitor concentrations. After that, plates were incubated for 72 hours before the addition of real-time glow (RTG) following the recommendation of the manufacturer (Promega Biotech AB, Sweden) and incubated at 37°C/5%CO_2_. Read-outs were performed 2, 24, 48, and 72 hours after adding RTG (days 3 to 6 after washing) using a temperature-controlled Tecan plate reader (37°C; acquisition integration time: 1.0 s). The effect of the combinations was assessed using the Chou-Talalay combination index ^26^, the MuSyC algorithm ^27^, and the SynergyFinder Plus package ^28^.

## Results

### ^177^Lu-lilotomab satetraxetan has cytotoxic activity across different models of lymphomas and is more active in B-than T-cell lymphomas

^177^Lu-lilotomab satetraxetan presented a median IC50 of 5.31 μg/mL (95% CI, 4.42-8.32) across 55 cell lines derived from different lymphoma subtypes. Activity was stronger in B-cell than in T-cell lymphomas (P=0.003) (Table 1; Figure 1; Supplementary Table S1).

**Table 1.** Anti-tumor activity of ^177^Lu-lilotomab satetraxetan based on histology after 144 hours of exposure. Data in ABC-DLBCL has been previously presented in ^19^.

|  | Median IC<sub>50</sub> (µg/mL) | 95% Conf. Interval | Number of cell lines |
| --- | --- | --- | --- |
| B-cell lymphomas | 4.54 | 1.92-6.8 | 46 |
| T-cell lymphomas | 16.42 | 6.44-299.57 | 9 |
| **Histology** |  |  |  |
| ABC-DLBCL | 1.8 | 0.49-16.06 | 8 |
| GCB-DLBCL | 4.99 | 0.37-8.61 | 19 |
| MCL | 5.02 | 1.4-18.25 | 10 |
| MZL | 3.45 | 1.12-10.07 | 6 |
| PMBCL | 0.093 | n.d. | 1 |
| CLL | 15.69 | 8.73-22.65* | 2 |
| ALCL, ALK+ | 10.59 | 4.96-320.9* | 4 |
| CTCL | 32.8 | 16.42-713.6* | 4 |
| PTCL-NOS | 6.29 | n.d. | 1 |

DLBCL, diffuse large B-cell lymphoma; ABC-DLBCL, activated B-cell like diffuse large B-cell lymphoma; GCB-DLBCL, germinal center B-cell type diffuse large B-cell lymphoma; MCL, mantle cell lymphoma; ALCL, ALK+, ALK-positive anaplastic large cell lymphoma; MZL, marginal zone lymphoma; CTCL, cutaneous T cell lymphoma; CLL, chronic B-cell leukemia; PMBCL, primary mediastinal large B-cell lymphoma; PTCL-NOS, peripheral T cell lymphoma, not otherwise specified. \* Upper) confidence limit held at the minimum (maximum) of the sample.

**Figure 1.**
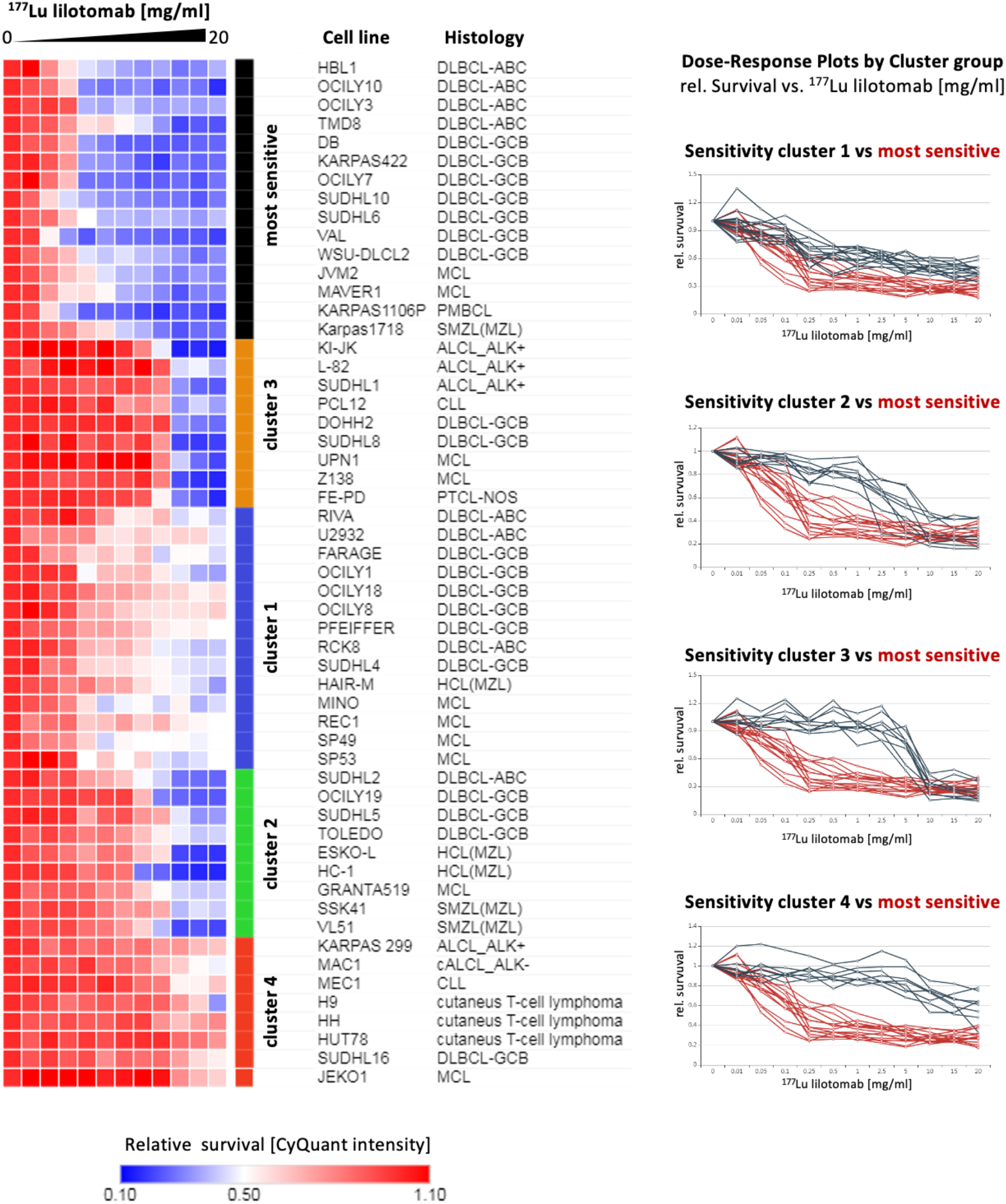
Supervised hierarchical clustering of lymphoma cell line panel by relative CyQuant signals six days after treatment with ^177^ Lu-lilotomab satetraxetan at doses ranging from 0.01 to 20 μg/mL. Signal intensities are normalized to untreated control and displayed as a color gradient [red to blue]

Unsupervised clustering of all the cell lines based on their dose-dependent response to ^177^Lu-lilotomab satetraxetan identified five groups of cell lines (Figure 1). One group included 16 cell lines with an IC50 below one μg/mL, derived only from B-cell lymphomas (75% diffuse large B-cell lymphoma, DLBCL). Another group had eight sensitive cell lines with an IC50 between 2.5 and 5 μg/mL (six were B-cell lymphomas). Similarly, the intermediate group (IC50s between 5 and 10 μg/mL) comprised eight cell lines with six of B-cell origin. The last two groups were composed of resistant cell lines. One had 15 cell lines, all B-cell lymphomas, with IC50s higher than ten μg/mL but still showing a dose-dependent response. The fifth cluster included eight cell lines (five of T-cell origin) completely resistant to ^177^Lu-lilotomab satetraxetan. CD37 expression was not different among these five clusters.

Focusing on DLBCL cell lines, representing the vast majority of our models and the most common type of lymphoma in patients, IC_50_ values did not differ based on the cell of origin, the presence of *BCL2* translocation (cell lines with n=15; without n.=11), of *MYC* translocation alone (with, n=10; without, n.=16) or concomitant to *BCL2* translocation (with, n=7; without n.=19), or *TP53* status (inactive, n=16; inactive, n=7).

### The *in vitro* cytotoxic activity of ^177^Lu-lilotomab satetraxetan correlated with its target expression

Taking advantage of available surface protein expression and RNA data obtained in our laboratory on the same cell lines used in this study ^14,23,24^, we analyzed whether the anti-tumor activity of ^177^Lu-lilotomab satetraxetan was affected by the expression levels of its target. ^177^Lu-lilotomab satetraxetan IC_50_ values were negatively correlated with CD37 surface protein expression across 54 cell lines, including both B and T cell lymphoma models (r=-0.48, P=0.003) and also within the group of B-cell lymphomas (r=-0.36, P=0.015) (Figure 2A-B). Similarly, the IC50 values were also inversely correlated with CD37 RNA levels, measured in 51 B and T cell lymphoma cell lines via a microarray-based technology (Illumina HT-12 arrays)(Pearson correlation r=-0.4 P=0.003) (Figure 2C-D) and, although at lesser extent, when we considered only the B-cell lymphoma cell lines using a data set we recently obtained with RNA-Seq (r=-0.26 P=0.05) (Figure 2C).

**Figure 2.**
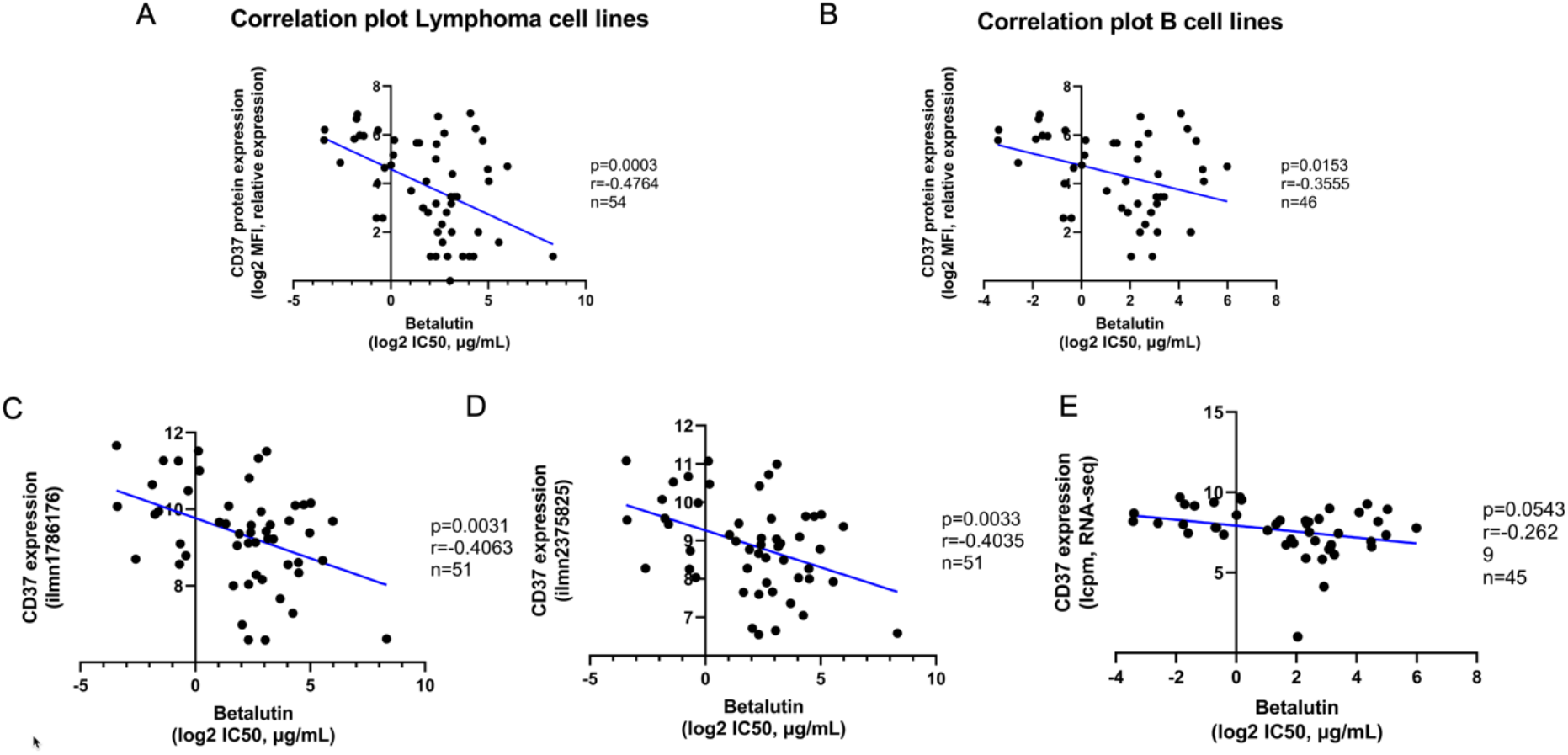
The *in vitro* cytotoxic activity of ^177^Lu-lilotomab satetraxetan correlated with CD37 expression. Pearson correlations between ^177^Lu-lilotomab satetraxetan activity, measured by IC_50_ values, with CD37 protein surface expression, measured by FACS in 54 B and T cell lymphomas (A) and or in 46 B-cell lymphomas (B) and with CD37 RNA levels, measured, by the two different probes on the Illumina HT-12 arrays, in 51 B and T cell lymphomas (C, D) and, via total RNA-Seq in 45 B-cell lymphomas (E).

Supplementary Figure S1 shows that B-cell lymphoma cell lines have higher CD37 expression levels than T-cell lymphomas (P<0.001), in agreement with the observed distribution of IC50 values.

### Transcriptome identifies genes associated with resistance to ^177^Lu-lilotomab satetraxetan and venetoclax as a novel active combination partner

To identify possible biomarkers of response and drugs that could be combined with ^177^Lu-lilotomab satetraxetan, we performed an exploratory analysis to compare the transcriptome of sensitive and resistant cells taking advantage of targeted RNA-seq profiling dataset (GSE94669 ^23^) we previously obtained in the same cell lines of the screening panel using a platform specifically designed to investigate potential therapeutic targets and drug response markers (HTG EdgeSeq Oncology Biomarker panel).

Due to sample size, we focused on DLBCL cell lines. Based on median IC50 values, GCB- and ABC-DLBCL cell lines were divided into two groups (“sensitive,” IC50s < 5 μg/mL; “resistant,” IC50s > 5 μg/mL), and a supervised analysis identified transcripts that were differentially expressed (fold change >|2|; P < 0.05) (Supplementary Table 2). In GCB-DLBCL cell lines, the transcripts up- and down-regulated in resistant cells were 125 and 85, respectively. In ABC-DLBCL, resistant cells had 95 up-regulated and 68-downregulated genes. Table 2 shows the seven differentially expressed genes, all up-regulated in resistance, and shared by both GCB- and ABC-DLBCL cells.

**Table 2.**
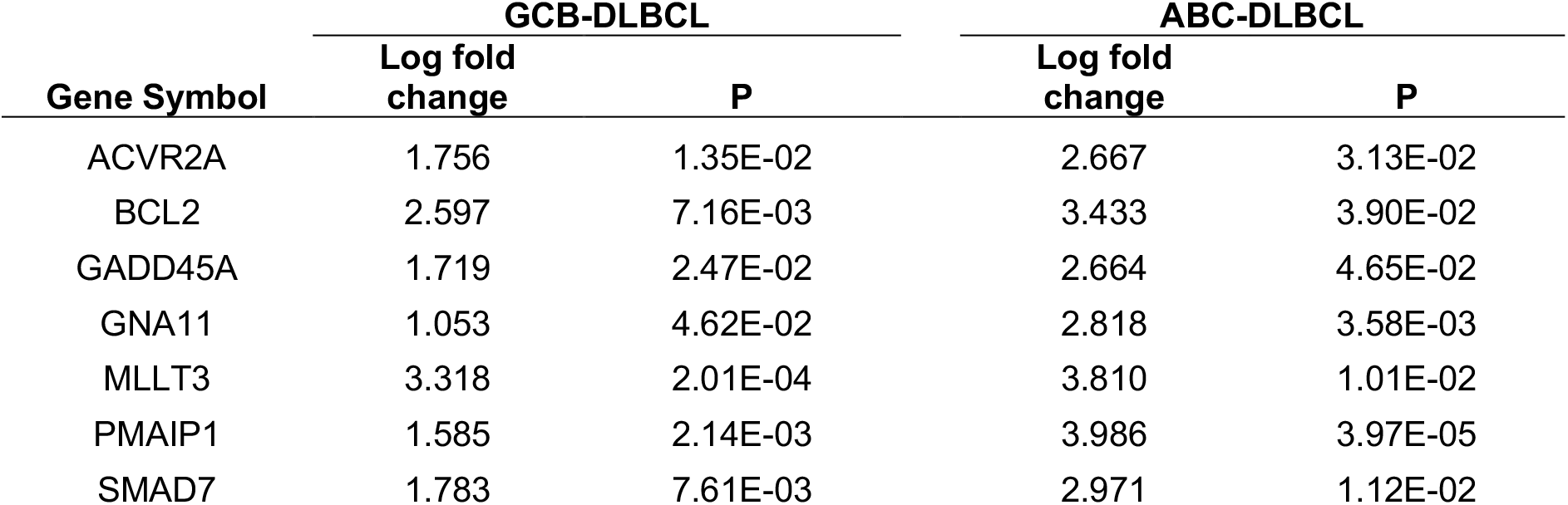
Genes were differentially expressed between resistant and sensitive cell lines in GCB- and ABC-DLBCL. ABC-DLBCL, activated B-cell like diffuse large B-cell lymphoma; GCB-DLBCL, germinal center B-cell type diffuse large B-cell lymphoma

Since the anti-apoptotic protein BCL2 was among the transcripts up-regulated in cell lines resistant to ^177^Lu-lilotomab satetraxetan, we assessed the combination of the radioimmunoconjugate with the BCL2 inhibitor venetoclax in two DLBCL cell lines, Toledo and DOHH2. Overall, the addition of venetoclax improved the activity of ^177^Lu-lilotomab satetraxetan in both cell lines. Based on different synergy models, the combination with venetoclax was mostly additive in Toledo, while in DOHH2, the effect of the BCL2 inhibitor was synergistic. According to the MuSyC algorithm, the addition of venetoclax increased the efficacy (i.e., the maximal effect) of ^177^Lu-lilotomab satetraxetan rather than the potency (i.e., lower dose with biological activity).

### ^177^Lu-lilotomab satetraxetan shows a different pattern of activity than the naratuximab emtansine and R-CHOP

To extend our findings, we compared the cytotoxic activity of ^177^Lu-lilotomab satetraxetan to what we have previously obtained using naratuximab emtansine (IMGN529), an ADC that incorporates the anti-CD37 humanized IgG1 monoclonal antibody K7153A conjugated to the maytansinoid DM1 as payload ^14^ and using the *in vitro* version of R-CHOP on the same cell lines ^14,29^.

In line with the differential expression of CD37 between B and T cell lymphomas, the IC50 values of ^177^Lu-lilotomab satetraxetan and naratuximab emtansine were positively correlated when considering both B and T cell lymphoma cell lines (r=0.28, P = 0.04) (Supplementary Figure S2A). However, when focusing on B-cell tumors, the two agents differed in their pattern of activity (r=0.13, P = 0.34) (Supplementary Figure S2B).

Finally, in DLBCL, IC_50_ values of ^177^Lu-lilotomab satetraxetan only partially correlated with what was obtained using R-CHOP (r=0.34, P = 0.085)(Supplementary Figure S3). Using the 75^th^ percentiles of the IC50 values in DLBCL (^177^Lu-lilotomab satetraxetan, 8.58 μg/mL; R-CHOP, 0.077 μg/mL), five DLBCL cell lines (TOLEDO, SU-DHL-16, PFEIFFER, U2932, RI-1), all characterized by inactivation of TP53, were resistant to both therapies. ^177^Lu-lilotomab satetraxetan maintained the activity in cell lines resistant to R-CHOP (SU-DHL-2, WSU-DLCL2), while R-CHOP was active in SU-DHL-5, OCI-LY-8, and OCI-LY-18, which were among the most resistant to the radioimmunoconjugate.

## Discussion

Here, we have assessed the anti-tumor effect of ^177^Lu-lilotomab satetraxetan in a panel of 55 cell lines derived from different lymphomas. The radioimmunoconjugate determined a dose-dependent anti-tumor activity across almost all cell lines, but we could identify subsets of cases with varying degrees of sensitivity. As expected, based on the CD37 expression pattern ^6-10^, T-cell lymphomas were less sensitive than B-cell lymphomas. This was also reflected in the observation that sensitivity to ^177^Lu-lilotomab satetraxetan was associated with the levels of its target when we considered the whole panel of cell lines and even when focusing to B-cell lymphomas only. This was similar to what was reported using the CD37-targeting ADC naratuximab emtansine ^7,14^. The analysis of clinical specimens collected in the clinical studies with ^177^Lu-lilotomab satetraxetan might confirm this observation using immunohistochemistry or RNA-based assays, even if the demonstration of a clear correlation between target expression and an antibody-based therapy is not straightforward in the clinical setting ^30^.

Focusing on B-cell lymphomas, we found differences in sensitivity based on histology or, within DLBCL, on the cell of origin. Furthermore, the anti-tumor activity of ^177^Lu-lilotomab satetraxetan was not negatively impacted by the presence of *BCL2* or *MYC* translocations (as individual or concomitant genetic events), but, moreover, the GCB-DLBCL cell lines more sensitive to the compound were also characterized by transcriptome similar to the double-hit gene expression signature recently reported associated with inferior outcome in patients with GCB-DLBCL ^31^.

We then looked for genes associated with sensitivity or resistance to ^177^Lu-lilotomab satetraxetan. Seven transcripts were more expressed in resistant than sensitive cells among GCB- and ABC-DLBCL cell lines. Some (ACVR2A, SMAD7, GADD45, and BCL2) likely played an active role in the sensitivity to ^177^Lu-lilotomab satetraxetan. ACVR2A, coding for the activin receptor type 2A, and SMAD7 are involved in TGFβ signaling ^32^, and, in a radiation-induced oral mucositis model, SMAD7 can protect cells from radiation, preventing apoptosis and sustaining cell proliferation ^33^, possibly playing a similar role also in lymphoma cells. Similarly, GADD45 protects hematopoietic cells from genotoxic stress ^34^. Finally, BCL2 is well known to block radiation-induced apoptosis ^35^. Importantly, when we combined ^177^Lu-lilotomab satetraxetan with the BCL2 inhibitor venetoclax, we observed an advantage in two DLBCL cell lines. The absence of a negative impact on sensitivity to ^177^Lu-lilotomab satetraxetan in cases where *BCL2* translocation, with or without MYC translocation, may seem contradictory to the association between BCL2 RNA levels and sensitivity. However, the role of BCL2 in the translocated cases is still unclear. BCL2 expression levels are not part of the double-hit gene expression signature associated with the presence of BCL2/MYC translocations ^31^. It is also noteworthy that while the presence of *BCL2* translocations may not be indicative, the overexpression of BCL2 could potentially predict higher responses to venetoclax ^36,37^.

We compared the activity of ^177^Lu-lilotomab satetraxetan with another CD37 targeting agent, the ADC naratuximab emtansine, which we had first studied in the same models ^14^. Across all the tested cell lines, ^177^Lu-lilotomab satetraxetan and naratuximab emtansine showed similar behavior, driven by CD37 expression on the cells’ surface. However, within the models derived from B-cell lymphoma, the two compounds differed, likely reflecting the different payloads and their mechanisms of action: the β-emitter and DNA-damaging ^177^Lu for the radioimmunoconjugate and the internalization-depending tubulin-targeting maytansinoid DM1 for the ADC. This suggests that approaches targeting the same surface markers are not necessarily cross-resistant.

Finally, we compared ^177^Lu-lilotomab satetraxetan with the DLBCL standard regimen, R-CHOP. Interestingly, although the activity of the two therapies differed across all the DLBCL cell lines, we identified a group of five cell lines resistant to both treatments. These cell lines were all characterized by TP53 inactivation, a feature known to play an essential role in the cancer cells’ response to radiation therapy, as reported in lymphoma cells ^38-40^. Conversely, there were cell lines that showed sensitivity to only one of the therapies, suggesting a lack of full cross-resistance and in agreement with the early signs of clinical activity with ^177^Lu-lilotomab satetraxetan in DLBCL patients after R-CHOP treatment ^21^.

In conclusion, in B-cell lymphomas, the CD37 targeting ^177^Lu-lilotomab satetraxetan presented preclinical anti-tumor activity, which was not cross-resistant with what was achieved with another CD37 targeting compound and with R-CHOP. The identification of transcripts, including BCL2, associated with resistance to ^177^Lu-lilotomab satetraxetan provided the rationale for adding the BCL2 inhibitor venetoclax, which increased the anti-lymphoma activity of the radioimmunoconjugate. Although the clinical development of ^177^Lu-lilotomab satetraxetan has now been put on hold, our data add to the safety profile, and the signs of activity originated in the early clinical trials and suggest that further clinical developments of the radioimmunoconjugate might be beneficial for lymphoma patients.

## Acknowledgements

This project was partially supported by research funds from Nordic Nanovector ASA. NM was supported by a Ph.D. Fellowship of the NCCR RNA & Disease, a National Centre of Competence in Research funded by the Swiss National Science Foundation (grant numbers 182880, 205601).

## Author Contributions

SP, JD: co-designed the study, supervised the study, interpreted data, and co-wrote the manuscript.

KBM, RG, AHVR-L: performed experiments and interpreted data.

NM, LC: performed data mining.

AJA: performed data mining and interpreted data. EG: interpreted data.

FB: co-designed the study, interpreted data, supervised the study, and co-wrote the manuscript. All authors reviewed and accepted the final version of the manuscript.

## Conflict of interests

AHVR-L JD: Nordic Nanovector ASA, employment and equity ownership. KBM, RG, SP: Nordic Nanovector ASA, employment. AJA: travel grant from Floratek Pharma SA. LC: travel grant from HTG Molecular Diagnostics. FB: ADC Therapeutics, Bayer AG, BeiGene, Floratek Pharma, Helsinn, HTG Molecular Diagnostics, Ideogen AG, Idorsia Pharmaceuticals Ltd., Immagene, ImmunoGen, Menarini Ricerche, Nordic Nanovector ASA, Oncternal Therapeutics, Spexis AG; consultancy fee from BIMINI Biotech, Helsinn, Menarini; advisory board fees to institution from Novartis; expert statements provided to HTG Molecular Diagnostics; travel grants from Amgen, Astra Zeneca, Beigene, iOnctura

