## Supplementary tables and figures for "Comprehensive Analysis of ^177^Lu-lilotomab Satetraxetan in Lymphoma Cell Lines: Implications for Precision Radioimmunotherapy and Combination Schemes"

[1] Research & Development, Nordic Nanovector ASA, Oslo, Norway; [2] Dept Radiation Biology, Institute for Cancer Research, OUH Norwegian Radium Hospital, Oslo, Norway; [3] Institute of Oncology Research, Faculty of Biomedical Sciences, USI, Bellinzona, Switzerland; [4] SIB Swiss Institute of Bioinformatics, Lausanne, Switzerland; [5] Oncology Institute of Southern Switzerland, Bellinzona, Switzerland.

***Co-corresponding authors:**
-Prof. Francesco Bertoni, Institute of Oncology Research, via Francesco Chiesa 5, 6500 Bellinzona, Switzerland. Phone: +41 58 666 7206;

### Supplementary Figures and Tables

**Figure S1. Distribution of CD37 expression between B- and T-cell derived lymphoma cell lines.** A) CD37 surface expression between B- and T-cell lymphoma cell lines. B-C) CD37 RNA expression values measured via microarray between B- and T-cell lymphoma cell lines. ****, P<0.0001 as determined by the Mann-Whitney test.

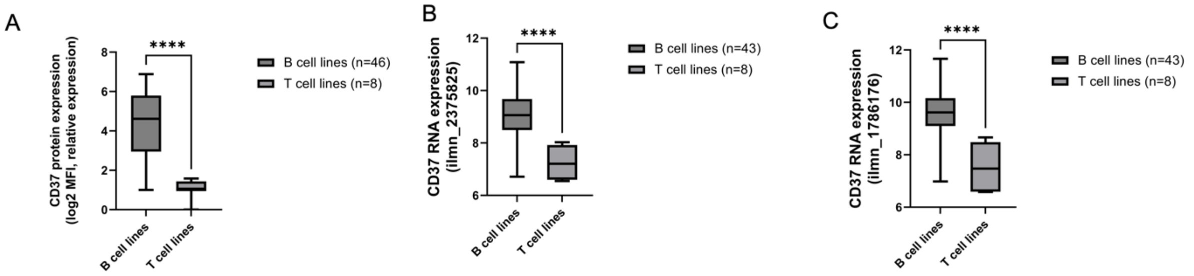


**Figure S2. Correlation between in vitro cytotoxic activities of $^{177}$Lu-lilotomab satetraxetan and the anti-CD37 ADC naratuximab emtansine in cell lines derived from 54 B- and T- cell lymphomas (A) and in 46 derived only from B-cell lymphomas (B).** Pearson correlations between log2 IC50 of the two agents. Data obtained with naratuximab emtansine were previously presented [1].

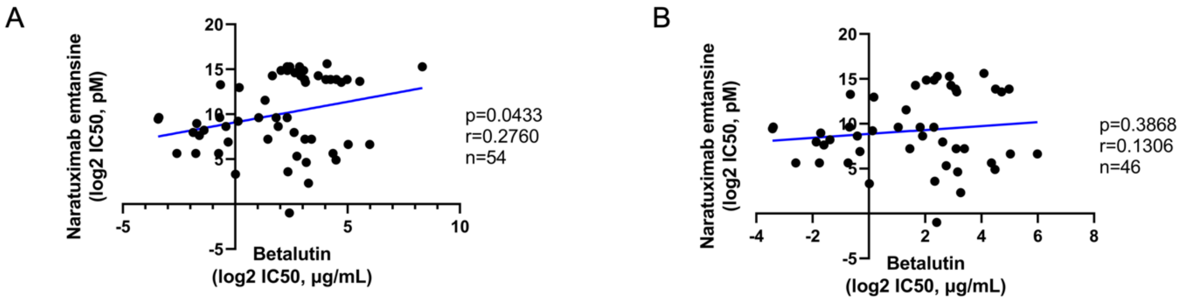

**Figure S3. Correlation between the activity of $^{177}$Lu-lilotomab satetraxetan and R-CHOP in 27 DLBCL cell lines.** Pearson correlations between log2 IC50 obtained with $^{177}$Lu-lilotomab satetraxetan or with R-CHOP. Data obtained with R-CHOP were previously presented [2].

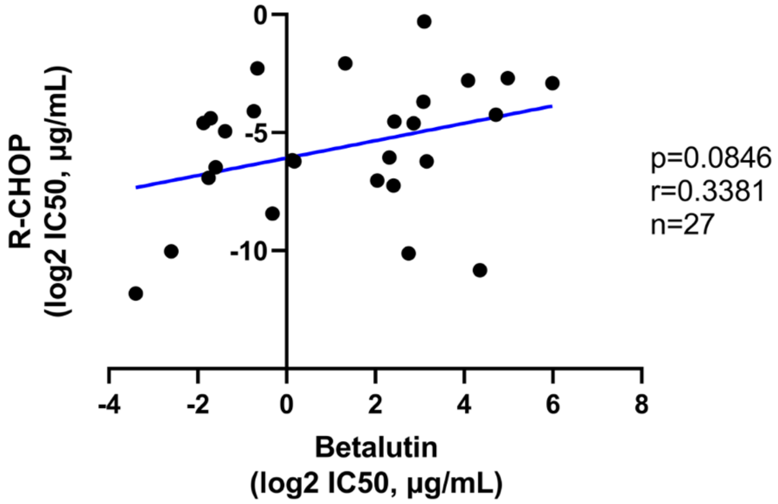

**Supplementary Table 1. Anti-tumor activity of $^{177}$Lu-lilotomab satetraxetan based in lymphoma cell lines after 144 hours of exposure.** Data in ABC-DLBCL has been previously presented [3].

| Cell line | Histology | IC50 (µg/ml) | Inactive TP53 | BCL2 transloc. | MYC transloc. | MYC + BCL2 transloc. |
|---|---|---|---|---|---|---|
| DB | GCB-DLBCL | 0.305 | yes | yes | no | no |
| DOHH2 | GCB-DLBCL | 4.985 | no | yes | yes | yes |
| ESKOL | MZL | 3.760 | . | . | no | . |
| FARAGE | GCB-DLBCL | 6.722 | yes | no | no | no |
| FE-PD | PTCL-NOS | 6.287 | . | . | . | . |
| GRANTA519 | MCL | 3.532 | yes | no | no | no |
| H9 | SS | 16.420 | . | . | . | . |
| HAIR-M | MZL | 6.158 | . | . | no | . |
| HBL-1 | ABC-DLBCL | 0.599 | yes | no | no | no |
| HC1 | MZL | 2.059 | . | . | no | . |
| HH | CTCL | 713.600 | . | . | . | . |
| HUT-78 | SS | 46.590 | . | . | . | . |
| JEKO1 | MCL | 22.400 | yes | no | no | no |
| JVM2 | MCL | 0.752 | no | no | no | no |
| KARPAS 1106P | PMBCL | 0.093 | . | . | no | . |
| KARPAS 1718 | MZL | 1.011 | yes | . | no | . |
| KARPAS 299 | ALCL | 320.900 | yes | . | . | . |
| KARPAS 422 | GCB-DLBCL | 0.383 | yes | yes | no | no |
| KI-JK | ALCL | 4.960 | . | . | . | . |
| L-82 | ALCL | 12.930 | yes | . | . | . |
| MAC1 | ALCL | 19.020 | . | . | . | . |
| MAVER1 | MCL | 0.621 | yes | . | yes | . |
| MEC1 | CLL | 22.650 | yes | . | no | . |
| MINO | MCL | 2.746 | yes | . | yes | . |
| OCI-LY-1 | GCB-DLBCL | 5.313 | yes | yes | no | no |
| OCI-LY-18 | GCB-DLBCL | 26.350 | yes | yes | yes | yes |
| OCI-LY-19 | GCB-DLBCL | 4.117 | no | yes | yes | yes |
| OCI-LY-3 | ABC-DLBCL | 1.096 | no | no | no | no |
| OCI-LY-7 | GCB-DLBCL | 0.296 | yes | no | yes | no |
| OCI-LY-8 | GCB-DLBCL | 20.500 | yes | yes | yes | yes |
| OCILY10 | ABC-DLBCL | 0.274 | no | no | no | no |
| PCL-12 | CLL | 8.728 | . | . | no | . |
| PFEIFFER | GCB-DLBCL | 17.040 | yes | yes | no | no |
| RCK8 | GCB-DLBCL | 5.383 | no | no | no | no |
| REC1 | MCL | 32.620 | yes | . | no | . |
| RI-1 | ABC-DLBCL | 8.593 | yes | no | yes | no |
| SP49 | MCL | 5.079 | . | . | no | . |
| SP53 | MCL | 4.969 | . | . | no | . |
| SSK41 | MZL | 10.510 | . | . | no | . |
| SU-DHL-1 | ALCL | 8.257 | yes | . | . | . |
| SU-DHL-10 | GCB-DLBCL | 0.165 | yes | yes | yes | yes |
| SU-DHL-16 | GCB-DLBCL | 63.490 | yes | yes | no | no |
| SU-DHL-2 | ABC-DLBCL | 2.506 | no | no | no | no |
| SU-DHL-4 | GCB-DLBCL | 0.330 | yes | yes | no | no |
| SU-DHL-5 | GCB-DLBCL | 8.931 | . | no | no | no |
| SU-DHL-6 | GCB-DLBCL | 0.804 | yes | yes | no | no |
| SUDHL8 | GCB-DLBCL | 7.270 | . | no | yes | no |
| TMD8 | ABC-DLBCL | 1.131 | no | no | no | no |
| TOLEDO | GCB-DLBCL | 8.476 | yes | yes | yes | yes |
| U2932 | ABC-DLBCL | 31.580 | yes | no | no | no |

| | | | | | | |
|---|---|---|---|---|---|---|
| UPN1 | MCL | 9.598 | yes | . | no | . |
| VAL | GCB-DLBCL | 0.095 | no | yes | yes | yes |
| VL51 | MZL | 3.148 | . | . | no | . |
| WSU-DLCL2 | GCB-DLBCL | 0.635 | yes | yes | no | no |
| Z138 | MCL | 7.534 | . | . | yes | . |

**Supplementary Table 2. Supervised analysis of resistant versus sensitive ABC- and GCB-DLBCL cell lines.** Limma analysis of the dataset GSE94669 [4] obtained with the HTG EdgeSeq Oncology Biomarker panel.

**Supplementary Table 3. Combinatorial effect of adding the BCL2 inhibitor venetoclax to $^{177}$Lu-lilotomab satetraxetan in two DLBCL cell lines.** Red: synergistic, orange: additive, green: antagonistic, black: no effect).

| Line (time) | Chou Talalay | MuSyc potency by drug1 | MuSyc potency by drug2 | MuSyc efficacy | ZIP score | HSA score | Bliss score | Loewe score |
|---|---|---|---|---|---|---|---|---|
| DOHH2 (day3) | 1.59 | -3.49 | -3.61 | 1.00 | 14.78 | 15.28 | 11.65 | 16.94 |
| DOHH2 (day4) | 7.35 | -3.59 | -3.73 | 1.06 | 7.00 | 12.14 | 1.33 | 13.22 |
| DOHH2 (day5) | 0.44 | 0.24 | -0.06 | 0.53 | 5.53 | 13.09 | 5.52 | 11.34 |
| DOHH2 (day6) | 2.54 | -3.68 | -3.87 | 0.78 | 12.07 | 11.38 | 11.06 | 11.46 |
| TOLEDO (day3) | 13.65 | 3.77 | -3.00 | 0.02 | 0.90 | 1.21 | -0.05 | 2.81 |
| TOLEDO (day4) | 0.64 | -3.79 | -4.04 | 0.19 | 0.78 | 4.26 | -0.08 | 4.82 |
| TOLEDO (day5) | 7.98 | -3.89 | -4.10 | 0.17 | 2.09 | 5.05 | 1.83 | 6.99 |
| TOLEDO (day6) | 0.47 | -3.71 | -3.97 | 0.48 | -1.98 | 7.61 | -0.33 | 8.70 |
